# Phasic arousal sharpens, rather than amplifies, goal-dependent feedback control

**DOI:** 10.64898/2026.07.02.736069

**Authors:** Clara Braconnier, Luca Creitz, Frédéric Crevecoeur, Julie Duque

## Abstract

Voluntary movements tolerate variability that does not threaten the goal: when reaching toward a wide target the hand can drift sideways, whereas a narrow target demands tighter control. The nervous system enforces this through corrective feedback responses that scale with task demands, yet how this scaling is set remains unclear. Arousal is a candidate, but how it would act is contested: the classical view casts it as a global amplifier that raises neural gain uniformly, whereas a more recent account holds that it is selective, strengthening task-relevant signals while suppressing others. These accounts make opposite predictions for goal-dependent feedback control, which we tested causally. In 27 human participants (24 ± 2.7 years old) reaching toward narrow or wide targets on a KINARM robot, a 4-second pulsed train of transcutaneous auricular vagus nerve stimulation (taVNS) evoked a phasic boost of arousal before each movement, confirmed by pupil dilation. On a subset of trials the robot applied a mechanical perturbation during the reach, and we measured the corrective muscle activity it evoked in the short- and long-latency reflex windows. The results favored the selective account. Phasic arousal sharpened corrections to the goal: it amplified the difference between narrow and wide responses in the long-latency window, the established readout of goal-dependent control, and suppressed the short-latency response, normally insensitive to the goal, specifically for the wide target, where deviations are harmless. Phasic arousal therefore does not energize the motor system indiscriminately but allocates corrective control to where it serves the task, extending the principle of selective gain, established in perception and memory, to the online control of movement.

## INTRODUCTION

Voluntary movements rely on continuous integration of delayed sensory feedback, enabling updates of online motor commands to achieve the task goal, despite sudden changes in our environment^1,2^. However, both sensory signals and motor commands are inherently noisy, introducing uncertainty in body state estimation and movement execution^3^. In addition, voluntary movement is inherently redundant, as the same behavioral goal can be achieved through many different motor commands and trajectories^4,5^.

A highly influential framework to explain how voluntary movements are controlled is Stochastic Optimal Control^4^ (SOC). Rather than selecting a single predetermined trajectory, SOC proposes that the nervous system implements a control policy, which adjusts motor commands online as a function of the evolving state of the body and the environment. These control policies are assumed to comply to a cost function which defines multiple objectives, including achieving the task goal while limiting the amount of motor effort or energy required^4,6^. As a result, only deviations threatening the task success are corrected, while variability is tolerated in dimensions that do not affect the movement goal. For instance, when reaching toward a wide target, movements can vary substantially along the target width without compromising success. In contrast, reaching toward a narrow target requires stricter control, as variability along this thin dimension increases the risk of missing it^7^.

A powerful way to probe control policies is to apply mechanical perturbations during reaching movements. The corrective muscular activity following a perturbation can be divided into an early short-latency (SL) reflex component, which originates primarily from spinal circuits, and a later long-latency (LL) response, which involves both spinal and supraspinal structures^8,9^. Importantly, LL responses are strongly modulated by the control policy: they are typically larger when reaching toward narrow targets and smaller when reaching toward wide targets^10^. Current evidence points to a distributed network involving sensorimotor cortex, cerebellum, and spinal circuits that integrate descending signals with sensory feedback^11,12^. However, how this network specifically regulates corrective responses in relation to a given control policy remains an open question. One plausible candidate that has been almost entirely overlooked in this context is the ascending arousal system^13^.

Evidence from the animal literature indicates that arousal, in particular the locus coeruleus-norepinephrine (LC-NE) system, sets the neural gain of its target circuits^14^. Because the LC projects widely throughout the central nervous system^15^, including key motor nodes such as the sensorimotor cortex, cerebellum, and spinal cord^16,17^, phasic arousal could modulate neural gain within the circuits that implement reaching control. Classically, this gain modulation has been viewed as a broadly activating influence that uniformly raises the responsivity of LC targets; more recent accounts propose instead that it is selective, enhancing signals relevant to the current task while suppressing irrelevant ones^18–20^. These two views make opposing predictions for the effect of increased arousal on goal-directed feedback control. If arousal increases gain broadly, corrective responses to perturbations should strengthen similarly across targets, including the wide one, where corrections are not needed for success, leaving the contrast between narrow and wide conditions essentially unchanged. If instead it modulates gain selectively, as a function of task demands, it should act unevenly across conditions, mainly when accuracy is most at stake, and thereby sharpen this contrast.

To address this question, we combined a reaching task with a causal manipulation of arousal. Healthy participants performed reaching movements towards narrow or wide targets, with or without, mechanical perturbations, using a KINARM robot. Critically, immediately before movement initiation, short phasic increases in arousal were induced using transcutaneous auricular vagus nerve stimulation (taVNS). taVNS is a non-invasive technique in which electrodes placed on the outer ear stimulate the region innervated by the auricular branch of the vagus nerve, an exclusively afferent sensory pathway that engages the arousal circuitry^21–24^. Reaching kinematics and feedback responses to perturbations were compared between taVNS and a sham condition in which stimulation was applied to the earlobe, in separate blocks; pupillometry confirmed that taVNS reliably increased arousal. Strikingly, rather than the uniform facilitation expected of an activating system, taVNS selectively reduced corrective responses for the wide target, sharpening the contrast between narrow and wide conditions and favoring a selective, rather than broad, action of arousal on goal-directed feedback control. This suppression was, moreover, already evident in the short-latency reflex component, suggesting an influence as early as spinal circuitry.

## RESULTS

A total of thirty-five participants were recruited for this study. Twenty-seven participants completed the main experiment, while eight participated in the control experiment. In both experiments, participants performed right arm reaching movements from a start target toward either a narrow (square) or a wide (rectangle) goal target using a KINARM end-point robotic device (see Fig. 1A-C and Materials and methods for more details). On two-thirds of the trials, participants experienced a mechanical perturbation (MP) of 9 N applied randomly to the left or to the right. Participants were explicitly instructed to land anywhere within the goal target, despite the possible occurrence of a mechanical perturbation. Movement kinematics and EMG activity of the right Posterior Deltoid and the right Pectoralis Major were recorded and served as primary outcome measures. In the main experiment, before each reaching movement, an extended fixation period lasting 9–9.5 s was imposed, during which a 4-second train of either sham or taVNS stimulation was delivered to the ear in separate blocks (see Fig. 1C-E and Materials and methods for more details). To assess arousal activity during the task under taVNS or sham stimulation, pupil size, a well-established index of arousal^22,25^ was recorded continuously throughout each trial.

**Figure 1.**
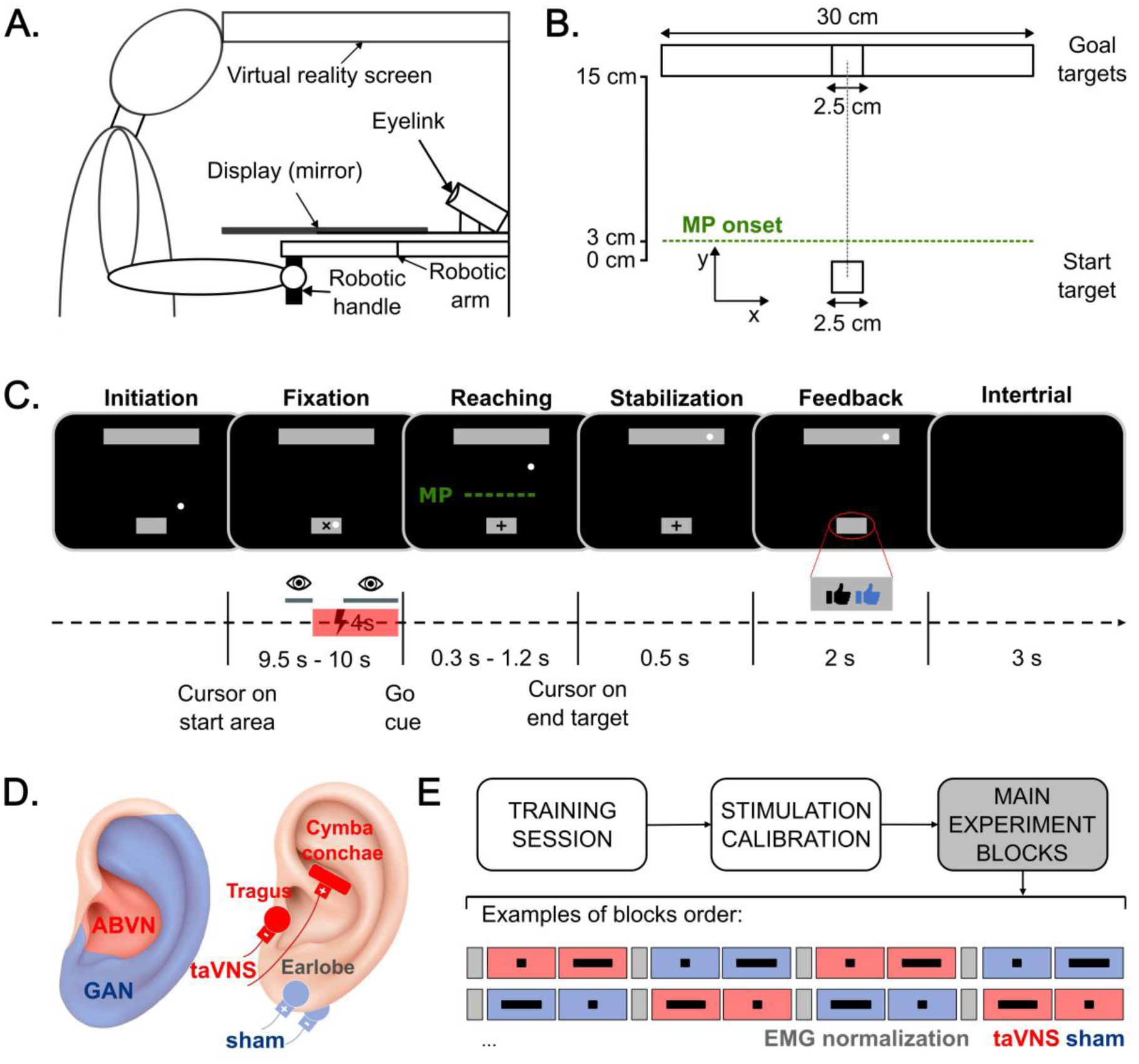
Study procedures. **A. Set-up and participant position.** Participants were seated at a KINARM end-point robotic device and performed right-arm reaching movements with the robotic handle while pupil size was recorded using an EyeLink eye tracker embedded in the system. The head was tilted ∼45° forward to ensure reliable eye tracking. Hand position was represented by a white cursor displayed via a mirrored virtual-reality setup occluding direct vision of the arm. **B. Scheme of the targets display.** Participants were asked to perform a reaching movement from a start target to a goal target, which was either narrow (square) or wide (rectangle). Targets are superposed for visualization purposes; only one goal target presented per trial. In 26 out of 39 trials per block, a 9N step force was applied to induce a mechanical perturbation (MP) either to the left or right (randomly). The green dashed line represents the y-position of the MP onset (here shown for visualization but not displayed on the screen). **C. Task sequence of events.** The sequence of events was identical for both goal target types (shown for wide target here). Each trial was initiated by placing the cursor within the start target, triggering a 9.5–10 s fixation period during which participants maintained their gaze on a fixation cross. After 5.5 s, a 4-s stimulation train (sham or taVNS, depending on the block) was delivered, followed by a random delay of 0–0.5 s. The fixation cross then changed into a “+” symbol, cueing participants to initiate a reaching movement toward the goal target within 0.3–1.2 s and to stabilize the cursor within the target for 0.5 s. The green dotted line (not displayed) represents the MP onset in perturbed trials, as in B. Feedback was provided for 2 s using a black thumb (stabilization success) and a blue thumb (movement speed success), followed by a 3-s intertrial interval. **D. transcutaneous auricular Vagus Nerve Stimulation (taVNS) set-up.** In the taVNS conditions, the electrodes were placed on the concha and the tragus of the left ear to stimulate the auricular branch of the vagus nerve (ABVN, in red). In the sham condition, the stimulation was applied to the earlobe, engaging the greater auricular nerve (GAN, in blue). **E. Experimental design.** The experiment began with a training session, followed by taVNS and sham stimulation calibration, and finally the main experiment blocks. Subjects completed 8 blocks of 39 trials (two blocks per conditions). The goal target type (narrow vs wide) alternated with each block, while the stimulation type (taVNS vs sham) alternated every two blocks. The starting experiment block was counterbalanced, resulting in 4 types of block sequences (two of them shown here). Additionally, before the experimental blocks and every two blocks thereafter, an EMG normalization block (grey rectangle) occurred.

### 1. EMG data

EMG signals were normalized (see Materials and methods for more details) and aligned to the moment the cursor crossed 3 cm beyond the start area, corresponding to perturbation onset. Then, mean EMG activities during unperturbed trials were subtracted to the EMG activity during perturbed trials. Analyses focused on reflex activity when the muscle acted as an agonist: leftward perturbations for the posterior deltoid, and rightward perturbations for the pectoralis major. Following established definition^26^, reflex activity was divided into two epochs: the short latency (SL) response (20–50 ms) and the long latency (LL) response (50–100 ms) after perturbation onset. Normalized EMG in these two epochs served as the primary reflex outcome measures: SL-reflex and LL-reflex.

Previous work grounded in optimal feedback control^7,10,27^ shows that long-latency reflex responses are shaped by task goals and are typically larger when accuracy demands are higher (e.g., narrow vs wide targets). We therefore examined whether reflex measures varied with goal target width under sham stimulation and whether this modulation was altered by taVNS stimulation. The lower panel of Figure 2A shows mean EMG traces for the posterior deltoid (leftward perturbation) under sham (left panel) and taVNS (right panel) for reaches toward wide and narrow targets. The upper panel of Figure 2A shows the corresponding SL- and LL-reflex values. Corresponding graphs are shown for the pectoralis major (rightward perturbation) on Suppl. Figure S.1

**Figure 2.**
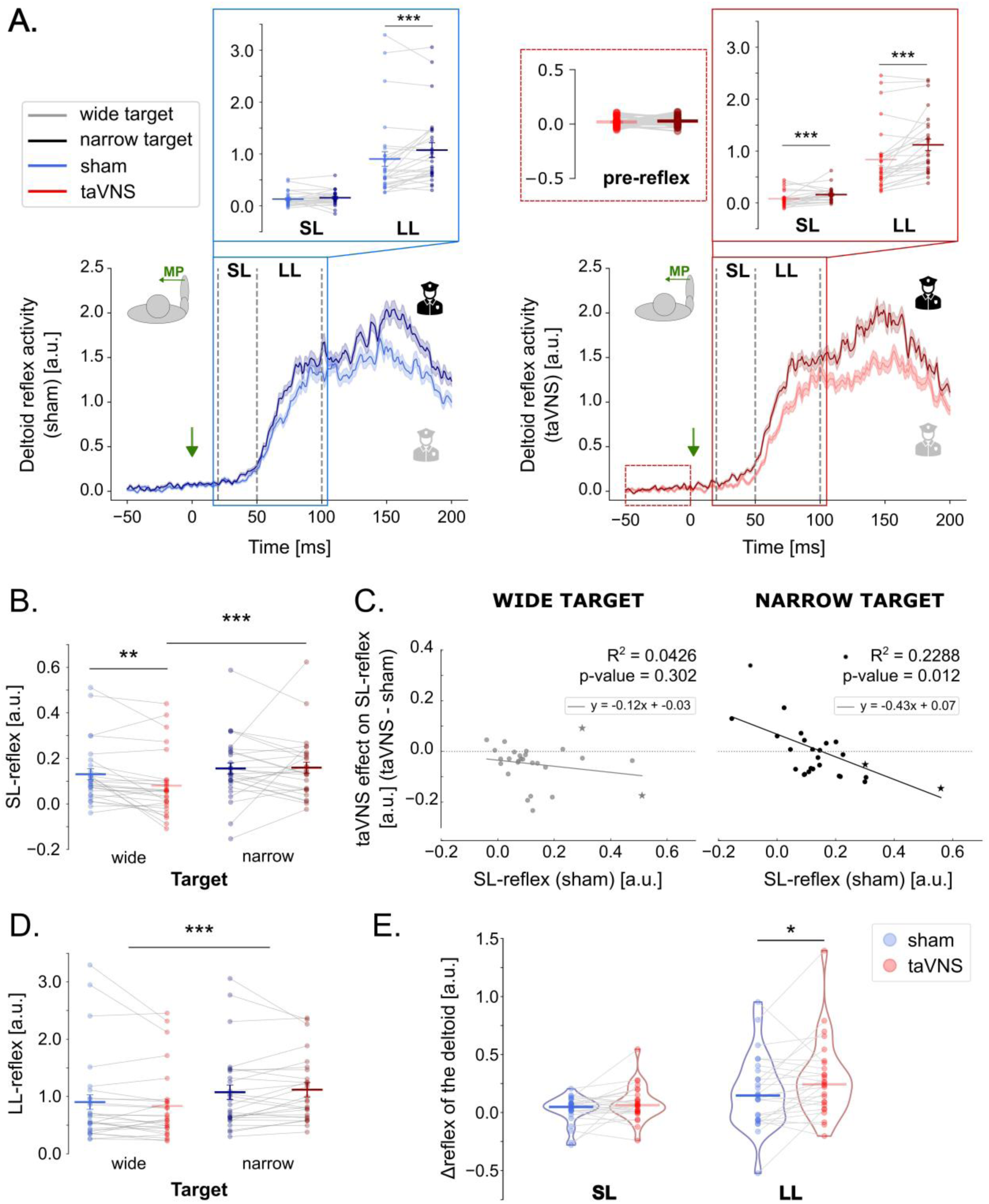
Posterior deltoid activity with leftward mechanical perturbation (MP). Data are shown for wide (light) and narrow (dark) targets under sham (blue) and taVNS (red) stimulation. **A. The lower panels show group-average EMG traces** (± SE) aligned to the mechanical perturbation onset and shown from −50 to 200 ms, under sham (left panel) and taVNS (right panel). Dashed vertical lines indicate the temporal boundaries of the short-latency (SL) and long-latency (LL) reflex epochs following the perturbation. Dashed box on the right panel corresponds to the temporal boundaries of pre-reflex. **The upper panels show SL and LL reflex components.** EMM (horizontal lines) and EMSD (vertical error bars) are represented with individual participant data displayed as dots connected with thin grey lines, shown as a function of Target-Width, under sham (left panel) and taVNS (right panel). On the right panel, the pre-reflex component is also displayed. **B. SL-reflex component.** EMM (horizontal lines) and EMSD (vertical error bars) are represented with individual participant data displayed as dots connected by thin grey lines, shown as a function of Target-Width and Stimulation-Type. **C. Baseline-dependent taVNS modulation of SL-reflex.** Correlation between taVNS-related changes in SL-reflex (y-axis) and SL-reflex under sham (x-axis) for wide (left panel) and narrow (right panel) target. Note the absence of significant correlation for the wide target, but the negative correlation for the narrow target: lower SL-reflex under sham led to a taVNS-related increase while larger SL-reflex under sham led to a taVNS-related decrease. The two participants exhibiting exceptionally large sham LL-reflex values are represented as stars instead of dots. **D. LL-reflex component.** EMM (horizontal lines) and EMSD (vertical error bars) are represented with individual participant data displayed as dots connected with grey thin lines, shown as a function of Target-Width and Stimulation-Type. **E.** Δ**SL and** Δ**LL reflex indices.** Data are displayed as violin plots with median values indicated by horizontal lines and individual participant data displayed as dots connected with thin grey lines. Note the significant taVNS-related increase in ΔLL-reflex compared to sham (Wilcoxon signed-rank test p = 0.025).

#### 1.1. Under sham stimulation, goal target width affected long-latency reflex activity of posterior deltoid but not pectoralis major

Under sham stimulation (Fig. 2A, left panel), linear mixed-effect analyses revealed a significant main effect of Target-Width on LL-reflex activity (F(1, 1371) = 22.2, p < 0.001), with larger responses for narrow than wide target reaches, consistent with the view that long-latency responses reflect goal-directed control policies during reaching. No such effect was observed for the SL-reflex (F(1, 1371) = 2.85, p = 0.092). The same analysis applied to the pectoralis major (rightward perturbation) revealed no effect of Target-Width on either reflex component under sham stimulation, including the long-latency response (both F(1, 1373) < 3.33, both p > 0.068; see Suppl. Fig. S.1, left panel). This absence of a goal target effect for the pectoralis major was unexpected and contrasted with previous findings. Notably, although present, the Target-Width effect was also relatively subtle for the posterior deltoid, compared to past studies using a comparable design. Yet a major difference here is that pupillometry required to implement an extended fixation period to obtain optimal pupil data, which might have promoted a center-aiming strategy across conditions, effectively reducing the functional distinction between narrow and wide targets. To test this possibility, we ran a control experiment on eight additional participants (4 women; mean age 24 ± 2 years old), who performed the same task with or without extended fixation in separate blocks (without any taVNS/sham stimulation), confirming attenuated target-related differences in the latter condition (see supplementary materials and Suppl. Fig. S.2).

#### 1.2. Under taVNS stimulation, goal target width affected short-latency and long-latency reflex activity of both posterior deltoid and pectoralis major

We then fitted the same linear mixed-effect model to the taVNS condition (Fig. 2A, right panel). Similar to the observation made under sham stimulation, we found a significant main effect of Target-Width on the LL-reflex (F(1, 1371) = 76.0, p < 0.001). Unexpectedly, here the model also revealed a Target-Width effect on the SL-reflex (F(1, 1371) = 29.1, p < 0.001). To exclude the possibility that the observed effect on the SL-reflex stemmed from gain scaling, potentially driven by higher EMG activity for the narrow target *prior* to the perturbation, we replicated the analysis on the mean EMG activity during the 50 ms pre-perturbation epoch (pre-reflex; Fig. 2A, right panel). Target-Width effect was not significant during this epoch (F(1, 1371) = 0.906, p = 0.341), suggesting that Target-Width effect on the SL-reflex under taVNS is unlikely to be attributable to distinct muscular activity between narrow and wide targets before the perturbation and the taVNS mostly shaped corrective feedback responses following the perturbation.

A similar Target-Width modulation was also observed for the pectoralis major (see Suppl. Fig. S.1, right panel), with significant effects evident for both SL-reflex (F(1, 1373) = 4.52, p = 0.034) and LL-reflex (F(1, 1373) = 10.5, p < 0.001), but not prior to the perturbation (pre-reflex; F(1, 1373) = 0.0474, p = 0.828). However, given the lack of target-related modulation in the pectoralis major under sham stimulation, subsequent analyses focused exclusively on reflex activity in the posterior deltoid.

#### 1.3. taVNS reduced short-latency reflexes during wide target reaches

Because we observed a Target-Width effect on the SL-reflex under taVNS (Fig. 2A, right panel), despite the typical absence of such modulation at short latency in prior work and in our sham condition (Fig. 2A, left panel), we further analyzed SL-reflex activity as a function of stimulation. Interestingly, SL-reflex activity varied with stimulation type, as supported by the linear mixed-effect model indicating a significant interaction between Target-Width and Stimulation-Type (F(1, 2768) = 6.31, p = 0.012; Fig. 2B). Post hoc analyses revealed that the taVNS effect was specific to wide-target reaches: SL-reflex was significantly reduced under taVNS relative to sham for the wide target (p = 0.005).

#### 1.4. taVNS modulated short-latency reflexes during narrow target reaches in way that depended on their magnitude under sham

We next tested whether the taVNS-induced change in SL-reflex (taVNS – sham) depended on short latency reflex magnitude under sham stimulation, which corresponds to the baseline activity of the muscle. For wide target reaches, this relationship was not significant (R^2^ = 0.0426, p = 0.302; see left panel of Fig. 2C), indicating a relatively uniform reduction in SL-reflex under taVNS across participants. In contrast, for narrow target reaches, a significant negative correlation was observed (R^2^ = 0.2288, p = 0.012; see Fig. 2C, right panel). To confirm that this relationship was not driven by mathematical dependence between the variables, we ran surrogate analyses (Supplementary Material 1. Statistical analysis), which confirmed this effect (Suppl. Fig. S.3A), suggesting that taVNS selectively decreased SL-reflexes in participants with larger SL-reflex responses under sham stimulation, while enhancing them in those with smaller SL-reflex responses under sham conditions.

Together, these results indicate that, at short latency, taVNS reduced SL-reflex activity during wide-target reaches, whereas during narrow-target reaches its effect was baseline-dependent, enhancing responses in individuals showing small ones under sham and attenuating them in individuals showing larger responses under sham stimulation.

#### 1.5. taVNS had no reliable impact on long-latency reflex activity during narrow and wide target reaches

LL-reflex responses were also analyzed as a function of the goal target width and the stimulation type. The linear mixed-effects model showed a significant interaction between Target-Width and Stimulation-Type on LL-reflex responses (F(1,2768) = 5.175, p = 0.023). Post hoc analyses confirmed a robust Target-Width effect (both p < 0.001), with larger LL-reflex responses for narrow than wide targets (see Fig. 2D). Besides the interaction between stimulation and target, when the analyses were restricted to each target independently, stimulation-related differences within each target width were absent: the effect of taVNS was marginal for the wide target (p = 0.098) and non-significant for the narrow target (p = 0.213). Thus, at the group level, taVNS neither selectively increased LL-reflex responses for narrow targets nor globally reduced responses for wide targets.

Consistent with the SL analysis, we also tested the relationship between the individual taVNS-induced change in LL-reflex (taVNS − sham) and LL-reflex amplitude under sham, reasoning that a baseline-dependent modulation might mask an effect at the group level. However, this relationship was not robust: the negative correlations observed for both targets were driven by two influential participants and did not survive their exclusion (both p > 0.45). These analyses are reported in the Supplementary Material (Suppl. Fig. S3B-C).

#### 1.6. taVNS amplified the goal target-width effect for LL- but not SL-reflexes

To further assess how taVNS-related boosts of arousal shaped corrective EMG responses, we computed for each participant, a difference index (Δ = narrow − wide) for the SL and LL epochs (ΔSL-reflex and ΔLL-reflex; see Fig. 2E), capturing the overall target width effect at each epoch under taVNS and sham. Given the selective decrease in SL responses for wide targets reported above, we expected ΔSL-reflex to increase under taVNS. Surprisingly, it did not differ between stimulation conditions (Wilcoxon signed-rank test p = 0.134), a lack of effect that may reflect the baseline-dependent modulation of the narrow-target SL-reflex (reported above and discussed below). In contrast, and despite the absence of a clear effect on the raw LL data, ΔLL-reflex was significantly larger under taVNS than sham (Wilcoxon signed-rank test p = 0.025), suggesting that taVNS amplified the goal-dependent adjustment of the motor response typically expressed in the long-latency reflex, presumably through small changes in the narrow- and wide-target responses that were individually non-significant but combined in the difference index.

#### 1.7. taVNS increased deltoid activity during unperturbed trials for the narrow target

Further analysis focused on the deltoid activity during unperturbed trials. No effect of taVNS on deltoid activity was observed in the timing corresponding to pre-, SL-, or LL-reflex windows. However, an effect emerged later in the movement. When looking at the mean deltoid activity from 100ms to 180ms after the hand crossed the threshold corresponding to the perturbation onset during the perturbed trials, which corresponds to the voluntary epoch usually described in the literature^10,26^, an interaction between Target-Width and Stimulation-Type was found (F(1, 2774) = 6.50, p = 0.011). Post-hoc analyses revealed that taVNS significantly increased the deltoid activity for the narrow target (p = 0.03), leading to a significant deltoid activity difference between the two targets under taVNS stimulation (p = 0.001).

### 2. KINEMATIC DATA

Hand trajectories for the two kinds of goal targets and across condition including or not a perturbation are depicted in Figure 3A. Hand trajectories are all aligned to the moment the cursor crossed 3 cm beyond the start target, corresponding to the virtual threshold used to trigger the perturbation in the trials with lateral step loads. Two main kinematic signals were extracted: lateral eccentricity and forward velocity (Fig. 3B, left and right panels, shown for unperturbed trials). Lateral eccentricity was defined as the deviation of the hand position along the x-axis, computed as the distance between the hand x-coordinate and the straight line connecting the center of the start target to the center of the goal target (grey dotted line in Fig. 3B). Forward velocity corresponded to the hand velocity along the y-axis. Both kinematic signals were aligned to the same reference event as the hand trajectories (represented as the green dotted line in both graphs) and were considered for a window extending from 200 ms before to 700 ms after this event. From these two signals, we extracted two measures of interest. First, we considered the absolute value of the lateral eccentricity at the end of the movement. This Eccentricity_Final_ therefore corresponds to the distance between the cursor and the goal target center at the end of the movement. Second, we extracted the peak value of the forward velocity, referred to as Velocity_Peak_. In the following sections, we examine how these kinematic measures varied with the goal target width and whether they were modulated by taVNS stimulation.

**Figure 3.**
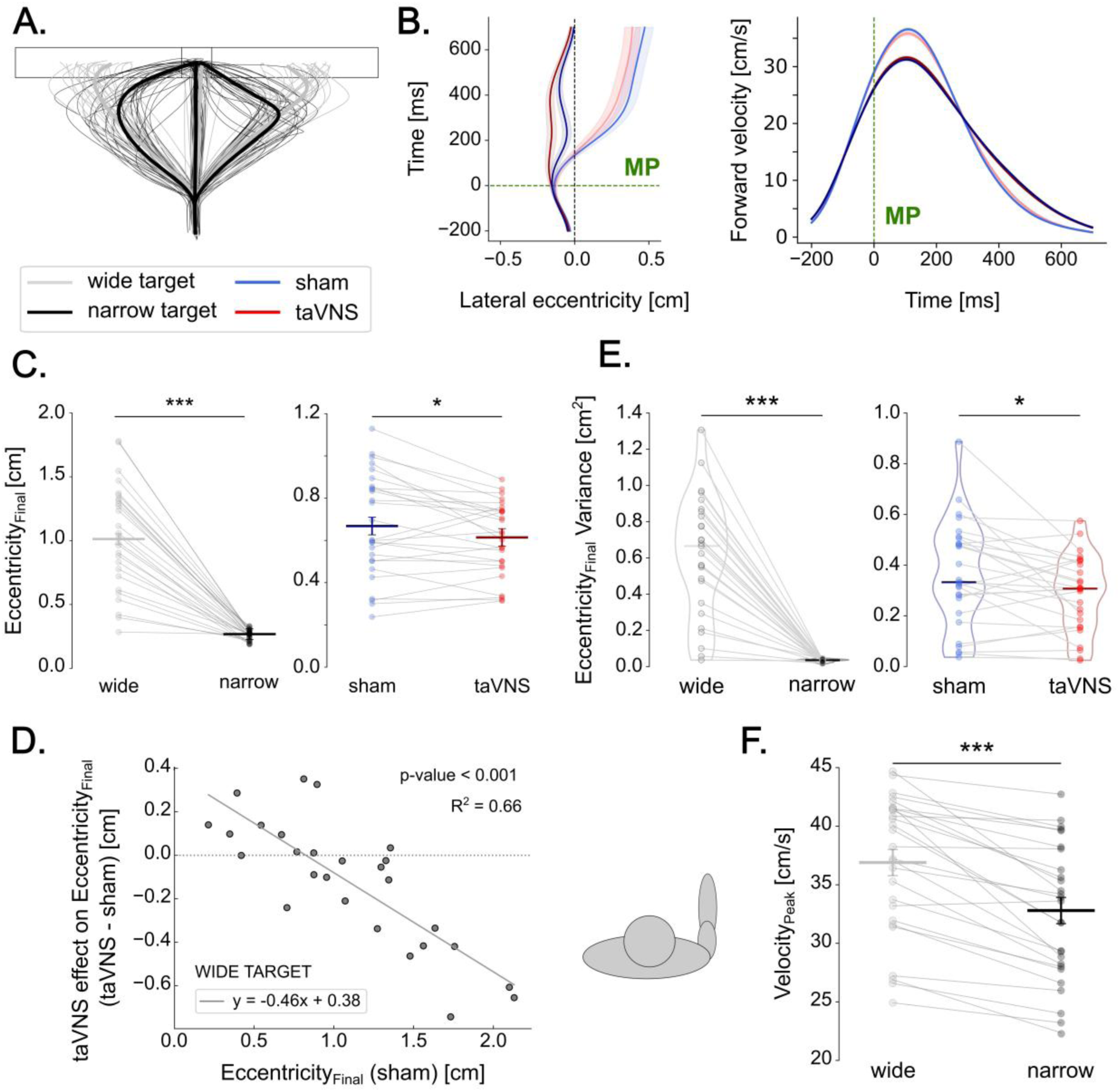
Kinematic data. Data are shown for wide (light) and narrow (dark) targets under sham (blue) and taVNS (red) stimulation. **A. Mean hand trajectories.** Thick traces show average hand trajectories across all participants, whereas lighter traces depict mean hand paths for individual participants. In each case, trajectories are shown for the three perturbation conditions (leftward, rightward, and no perturbation). Note that only panel A includes all three perturbation conditions; the remaining panels focus exclusively on trials without mechanical perturbation. **B. Kinematics signals.** Group mean (solid lines, ± SE) lateral eccentricity (left panel) and forward velocity (right panel) signals in unperturbed trials, aligned to the time point at which the mechanical perturbation (MP) would have occurred if the trial was perturbed (indicated by the green dotted line). Grey dotted line in the left panel connects the center of the start and goal targets. **C. Eccentricity_Final_.** EMM (horizontal lines) and EMSD (vertical error bars) are represented with individual participant data displayed as dots connected with thin grey lines. Left panel shows the effect of Target-Width, whereas right panel shows the effect of Stimulation-Type. **D. Baseline-dependent taVNS modulation of Eccentricity_Final_.** Negative correlation between the effect of taVNS on Eccentricity_Final_ (y-axis) and Eccentricity_Final_ in sham condition (x-axis) for the wide target. Note the taVNS-related increase in Eccentriciy_Final_ for individuals showing lower values under sham but the taVNS-related decrease in Eccentriciy_Final_ for individuals showing larger values under sham. **E. Variance of Eccentricity_Final_.** Data are displayed as violin plots with median values indicated by horizontal lines and individual data displayed as dots connected with thin grey lines. Left panel shows the effect of Target-Width, whereas right panel shows the effect of Stimulation-Type. **F. Velocity_Peak_.** EMM (horizontal lines) and EMSD (vertical error bars) are represented with individual data displayed as dots connected with thin grey lines, shown as a function of Target-Width. *: p < 0.5. ***: p < 0.001. EMM = estimated marginal means, EMSD = estimated marginal standard deviations.

#### 2.1. taVNS had no effects on kinematics data for perturbed trials

As in the literature, linear-mixed effects showed a main effect of Target-Width on Eccentricity_Final_ (F(1, 2190) = 2942.3, p < 0.001). Eccentricity_Final_ was significantly greater for the wide target (7.09 ± 0.43 cm) compared to the narrow target (0.354 ± 0.43 cm). However, no effects of Stimulation-Type (F(1, 2181) = 0.289, p =0. 591) nor interaction (F(1, 2181) = 0.056, p = 0.813) were observed. Moreover, for the Velocity_Peak_, only a main effect of Target-Width was observed (F(1, 1331) = 15.77, p < 0.001), with Velocity_Peak_ being lower for the narrow (40.3 ± 0.3 cm/s) than for the wide target (41.2 ± 0.3 cm/s).

Hence, although taVNS reliably modulated corrective EMG activity, this modulation did not produce a corresponding target-width-dependent change in movement kinematics. This dissociation points to a subtle influence of phasic arousal that influenced corrective EMG responses without reshaping the movement at the macroscopic level.

#### 2.2. taVNS shaped Eccentricity_Final_ in a goal-directed manner during unperturbed trials

As shown in Figure 3C, Eccentricity_Final_ varied with both the goal target width and the stimulation type. As such, Eccentricity_Final_ was larger when participants reached for the wide (1.01 ± 0.04 cm) compared to the narrow target (0.27 ± 0.04 cm), as supported by the linear mixed-effects model indicating a significant Target-Width effect (F(1,2596) = 1047.77, p < 0.001; Fig. 3C, left panel). Moreover, Eccentricity_Final_ was reduced under taVNS (0.61 ± 0.04 cm) compared to sham stimulation (0.67 ± 0.04 cm; Fig. 3C, right panel), as shown by a main effect of Stimulation-Type (F(1,2596) = 5.54, p = 0.019). These results indicate that while participants exploited the spatial redundancy offered by the wide target, a pattern consistent with previous findings, taVNS tended to constrain reaching endpoints toward the center, irrespective of the target width. Although the interaction between Target-Width and Stimulation-Type did not reach statistical significance (F(1, 2594) = 3.41, p = 0.065), its proximity to the conventional threshold prompted exploratory post-hoc comparisons. These analyses tentatively suggest a taVNS effect specific to the wide target (p = 0.016), with Eccentricity_Final_ decreasing from 1.06 ± 0.04 cm under sham stimulation to 0.96 ± 0.04 cm under taVNS (see Suppl. Fig. S.4A). No comparable effect was observed for the narrow target (p = 0.984).

Correlation analyses then tested whether the taVNS-induced change in Eccentricity_Final_ (taVNS - sham) related to the participants’ Eccentricity_Final_ under sham. A significant negative correlation was present for both targets (wide: R^2^ = 0.6606, p<0.001, Fig. 3D; narrow: R^2^ = 0.3905, p = 0.009, Suppl. Fig. S.4B). Surrogate analysis supported the correlation for the wide target only (Suppl. Fig. S4C), indicating that the narrow-target effect was attributable to mathematical coupling. A complementary surrogate analysis of the direction of the effect showed that, for the wide target, participants who under sham terminated near the target center showed a taVNS-induced increase in Eccentricity_Final_, whereas those who exploited more of the target’s spatial redundancy showed a taVNS-induced decrease. Consistent with this bidirectional convergence, inter-individual variability in Eccentricity_Final_ for the wide target was lower under taVNS than sham (Levene’s test: F = 5.550, p = 0.023; Fig. 3C, right panel).

Finally, we computed the intra-individual variance of Eccentricity_Final_ across trials within participants and the ANOVA revealed a significant main effect of both Target-Width (z = −10.79, p < 0.001) and Stimulation-Type (z = −2.45, p = 0.014). As shown in Figure 3E, the variance of Eccentricity_Final_ was significantly greater for the wide (0.616 ± 0.050 cm²) than for the narrow target (0.033 ±0.001 cm²), as previously reported in the literature^7^. More interestingly, taVNS generally reduced variance (0.287 ± 0.045 cm²) compared to sham (0.362 ± 0.060 cm²). The absence of interaction (z = 1.72, p = 0.085) suggests that this effect occurred regardless of the goal target width.

#### 2.3. taVNS did not impact Velocity_Peak_

The analysis of Velocity_Peak_ revealed a main effect of the Target-Width (F(1,2698) = 404.1, p < 0.001; see Fig. 3F), indicating that the peak velocity of reaching movements was significantly lower for the narrow (32.5 ± 0.01 cm/s) than for the wide target (37.0 ± 0.01 cm/s), as expected given the greater precision required to reach the former target. Neither the main effect of Stimulation-Type nor its interaction with Target-Width revealed any additional modulation of Velocity_Peak_ attributable to taVNS.

Overall, kinematic analyses of unperturbed trials indicate that taVNS biased movement endpoints by reducing their eccentricity, especially for the narrow target. However, the reduced endpoint eccentricity was conditional for the wide target: the direction and magnitude of the taVNS-induced shift depended on baseline performance under sham stimulation. In participants who showed greater endpoint eccentricity under the sham condition, taVNS shifted endpoints toward the target center. Conversely, in participants with lower sham eccentricity, taVNS moved endpoints away from the center. Overall, this resulted in a compression of the inter-individual distribution of endpoint locations compared to sham. Moreover, for both targets, taVNS significantly reduced intra-individual endpoint variability, indicating enhanced movement consistency around the new, taVNS-shifted endpoint. In contrast, movement velocity remained unaffected by taVNS.

### 3. Pupillometry data

Pupil size was monitored as an index of arousal, with the expectation that effective taVNS would elicit a pupil dilation relative to sham, consistent with previous reports^22,28–30^. Pupil size data were converted in z-scores computed across all conditions for each participant and aligned to the sham or taVNS stimulation onset. These pupil z-scores were then normalized relative to the mean z-score obtained during the last second of the inter-trial interval, when no target was displayed; this value was subtracted from z-scores obtained throughout the trial. From these signals, we first computed the Pupil_Baseline_, defined as the mean pupil size during a 1-s window preceding the stimulation onset (see Fig. 1C and 4AB). This measure served to evaluate baseline pupil size at trial onset (before stimulation) but when the start and goal targets were already displayed. Then, to allow for comparisons between conditions and participants, we subtracted this Pupil_Baseline_ from the remaining signal to obtain a relative measure of pupil dilation starting from stimulation onset (see Fig. 4, upper panel). From this signal, we selected a 2-s window, corresponding to the 2 last seconds of stimulation (see Fig. 1C and 4B). This Pupil_Stimulation_ measure reflects the pupil dilation elicited during taVNS or sham stimulation. The pupil analyses were restricted to unperturbed trials and to trials with leftward perturbations, corresponding respectively to those used for kinematic and deltoid reflex analyses.

**Figure 4.**
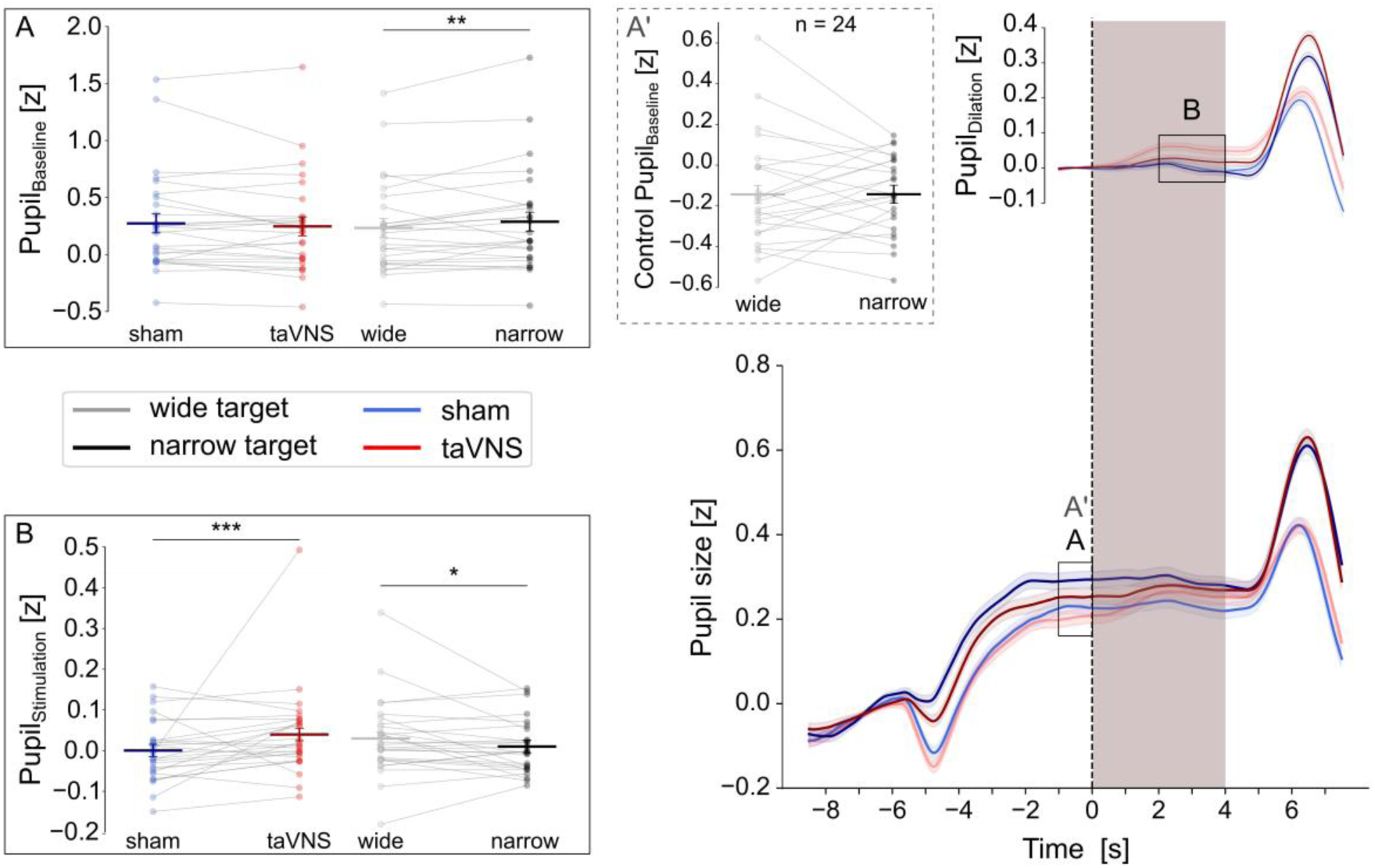
Pupillometry data. Data are shown for wide (light) and narrow (dark) targets under sham (blue) and taVNS (red) stimulation (shaded grey area). **Group-average pupil size traces.** Mean (± SE) traces are aligned to stimulation onset (dashed vertical lines). The lower panel shows pupil size for the whole trial duration, with the rectangle (AA’) indicating the window used to compute Pupil_Baseline_. The upper panel shows baseline-corrected pupil size from 1 second before stimulation onset, with the rectangle (B) indicating the window used to compute Pupil_Dilation_. **A. Pupil_Baseline_.** EMM (horizontal lines) and EMSD (vertical error bars) are represented with individual participant data displayed as dots connected with thin grey lines, as a function of Target-Width and Stimulation-Type. **A’. Control Pupil_Baseline_.** Mean pupil size over the same time window (AA’) as is A in a separate, unpublished follow-up experiment on 24 healthy participants using a similar paradigm and for which the targets were matched for luminance. Mean pupil size EMM (horizontal lines) and EMSD (vertical error bars) are represented with individual participant data displayed as dots connected with thin grey lines, as a function of Target-Width and Stimulation-Type. **B. Pupil_Stimulation_.** Same data organization as in A. *: p < 0.5. **: p < 0.01. ***: p < 0.001. EMM = estimated marginal means, EMSD = estimated marginal standard deviations.

#### 3.1. taVNS had no impact on Pupil_Baseline_

Neither Stimulation-Type nor the Perturbation-Type (no perturbation vs. leftward perturbation) affected Pupil_Baseline_ and no interactions were significant. This was expected since both (stimulation and perturbation) occurred after the window used to compute the baseline. The analysis did, however, reveal a significant main effect of Target-Width (F(1, 4872) = 8.253, p = 0.004) : baseline pupil size was larger during display of the narrow target (0.288 ± 0.0832 z) than the wide target (0.233 ± 0.0833 z). This could reflect a difference in arousal; the narrow target, being harder to reach, may elevate arousal even before stimulation and movement onset. Alternatively, it could be luminance-driven, since the smaller narrow target has lower overall luminance, which typically increases pupil size.

To distinguish between these possibilities, we examined baseline pupil size over the same time window in a separate, as-yet-unpublished follow-up experiment that used a similar reaching paradigm (movements toward a narrow or wide target) but in which the two targets were matched for luminance (n = 24; 20 women; mean age 24 ± 2 years; Fig. 4A’). In this dataset the Target-Width effect was abolished (F(1,6714) = 8.9 × 10^−4^, p = 0.976), indicating that the baseline pupil difference in the present study most likely reflects the luminance difference between targets rather than a difference in arousal.

#### 3.2. taVNS increased Pupil_Stimulation_

The analysis of Pupil_Stimulation_ revealed significant main effects of both Target-Width (F(1, 4870) = 6.13, p = 0.013) and Stimulation-Type (F(1, 4870) = 15.17, p < 0.001). Specifically, opposite to what was found for Pupil_Baseline_, Pupil_Stimulation_ was larger during reaches toward wide (0.032 ± 0.015 z) than narrow targets (0.002 ± 0.015 z). This difference may reflect the larger Pupil_Baseline_ observed for the narrow target, resulting in a smaller pupil response after baseline correction compared to the wide target. More interestingly, taVNS elicited greater Pupil_Stimulation_ (0.041 ± 0.016 z) than sham stimulation (−0.007 ± 0.015), indicating that taVNS effectively increased arousal. Importantly, this taVNS effect did not depend on the Target-Width (Interaction F(1, 4870) = 2.77, p = 0.096). Finally, Pupil_Stimulation_ was not affected by the presence of a perturbation (Perturbation-Type F(1, 4869) = 0.001, p = 0.921) and none of the other interactions reached significance. Taken together, these results indicate a global increase in pupil dilation, consistent with increased arousal under taVNS.

## DISCUSSION

We asked whether phasic arousal modulates online feedback control as a uniform increase in neural gain or as a selective, goal-dependent one. Across both reflex epochs, the modulation proved selective. Rather than strengthening corrective responses indiscriminately, taVNS reshaped them according to the goal of the movement: it sharpened the contrast between narrow and wide targets in the long-latency response, the component classically read as a signature of the control policy^9,12^, and, more strikingly, did so in part by reducing the short-latency correction precisely for the wide target, where deviations along the redundant dimension are tolerated. The convergence of these two effects is difficult to reconcile with a broadly activating role of the arousal system^31,32^ and instead provides a behavioral instance of selective modulation of corrective responses, that is, the enhancement of task-relevant signals together with the relative suppression of task-irrelevant ones^18,19^. Whereas selective gain has been characterized largely in perception, attention, and memory, our findings extend it to the online control of movement, showing that arousal can scale sensorimotor feedback to the task goal rather than uniformly energizing it. That such structure was already expressed at short latency, in a response generally automatic and goal-invariant^33^, is especially notable, although this early effect warrants confirmation given its novelty.

The pupillary data confirms that our manipulation engaged the arousal system. taVNS increased pupil size relative to sham during the last seconds of stimulation, but did not change baseline pupil size, taken before stimulation onset, indicating that the effect did not carry over between trials. taVNS thus produced transient, phasic boosts of arousal rather than a sustained shift in tonic state^14,28,30^. Which neuromodulatory pathway carried this effect cannot be settled here. Stimulation of the auricular branch of the vagus nerve activates the nucleus tractus solitarius, which projects to the locus coeruleus^34–36^, repeatedly reported among the strongest responders to taVNS, making an LC-NE contribution the most parsimonious account. The same afferent, however, may also engage other neuromodulatory systems, directly or via relays, including serotonergic, cholinergic, and dopaminergic ones, so a purely noradrenergic interpretation cannot be asserted^35,37–39^. What the pupil results establish unambiguously is that taVNS engaged the arousal system phasically; the selective signature described above must therefore be read as an effect of phasic arousal, whatever its precise neurochemical substrate.

The long-latency results provide the clearest evidence that arousal scaled corrective responses to task demands rather than raising them uniformly. Under sham, long-latency responses were larger for narrow than for wide targets, the expected signature of a goal-dependent control policy^7,10^. taVNS amplified this differentiation: the difference between the narrow and wide responses in the long-latency epoch was significantly larger under taVNS than under sham. This amplification did not arise from a global increase in neural gain, since neither the narrow nor the wide response shifted reliably on its own, but from a sharpening of the contrast between them. A uniform boost of neural gain would have raised responses across targets and, if anything, compressed this contrast; the opposite was observed. The effect is therefore more naturally read as arousal tuning corrective responses to the goal than as an indiscriminate strengthening of them. Because it is carried by the difference index rather than by either target alone, it should be taken as evidence for sharpened goal-dependent modulation rather than for a change in any single corrective response.

The short-latency results extend this picture to a level where it was not expected. Goal-dependent modulation of upper limb short-latency responses is normally absent in this kind of task, or emerging only after extensive training^40^. Our sham condition reproduced this: it showed no reliable target effect at short latency. Under taVNS, by contrast, a clear target effect appeared, with larger responses for narrow than wide targets. This did not reflect a difference in pre-perturbation muscle activity, as pre-reflex in the 50 ms before perturbation did not differ between targets, ruling out background-dependent gain scaling. The effect was driven by a selective reduction of the short-latency response for the wide target under taVNS, a suppression of the very correction that is unnecessary when deviations along the target’s redundant dimension do not threaten success of the reaching movement. This goal-dependent pattern at short latency under taVNS was not confined to the posterior deltoid: as reported in the supplementary material, the pectoralis major, the agonist for perturbations in the opposite direction, likewise showed smaller short-latency responses for the wide than the narrow target under taVNS, and neither muscle showed such a difference under sham. This suppression did not, however, translate into a reliable change in the overall narrow–wide difference at short latency in the deltoid, because the narrow-target response was itself heterogeneous across participants, a dependence on behavior under sham we return to below. Critically, target width was held constant within a block, so participants had sustained knowledge of the goal under both stimulation conditions. The fact that this goal-related modulation emerged at short latency only under taVNS shows that goal knowledge alone did not produce it; arousal was required for goal information to shape the earliest, normally goal-invariant component of the corrective response.

Because the taVNS input was identical for narrow and wide targets, the dependence of the effect on target width cannot have been carried by the stimulation itself; it must instead arise from the interaction of arousal with goal-related signals. And because the short-latency loop is too fast to incorporate the perturbation through transcortical processing, those signals cannot have been engaged online but must instead have been set before the perturbation, biasing spinal processing in a goal-dependent way. This anticipatory reading is reinforced by the timing of the manipulation: taVNS was delivered before movement onset and was not time-locked to the perturbation, so the arousal it evoked acted as a state established in advance rather than as a response to the perturbation. Nor could this preparation have been keyed to a specific perturbation: a perturbation occurred on only two-thirds of trials and, when it did, was equally likely to be leftward or rightward, engaging different agonist muscles, with the same probabilities for narrow and wide targets. The goal-dependent setting of corrective responses therefore reflected a standing, direction-general readiness, scaled by whether a correction would matter for the goal rather than by any expectation of a particular perturbation. It also produced no goal-dependent change in the tonic motor state: the pre-reflex, as noted, carried no target difference, so the modulation acted on the reflex response itself rather than on the ongoing drive against which the perturbation arrived.

The main candidate pathway to reduce short-latency responses is through pre-synaptic inhibition engaged by taVNS. It has been shown that this inhibitory mechanism, unlike post-synaptic inhibition, can regulate spinal feedback by amplifying or suppressing proprioceptive signals conveyed to the motoneurons, allowing smooth movements^41–43^. This interpretation is further supported by the absence of direct effect of taVNS on long-latency responses. It rules out the possibility of an inhibition driven by gamma motoneurons, which would uniformly affect both response components by adjusting muscle spindle sensitivity^44,45^. Pre-synaptic inhibition could be mediated either via supra-spinal structures or through direct projections of taVNS to the spinal cord. However, the latter appears unlikely, as the observed modulation was goal-dependent, and prior evidence demonstrates that both norepinephrine and serotonin, two neuromodulators targeted by taVNS, consistently enhance motoneuron activity^46–49^. Independently of the mechanism of inhibition, what our data established is that goal-related structure reached the short-latency reflex only under arousal. We cannot tell whether arousal amplified a goal-dependent gating that is normally present but too weak to surface at this level, or enabled one that does not otherwise reach it; in either case, phasic arousal was the condition under which the goal came to shape spinal processing. Functionally, this amounts to a suppression of reflexive corrections that are irrelevant or counterproductive to the task, a capacity whose failure is illustrated by motor-impaired populations in whom excessive perturbation-evoked activity reflects an inability to suppress maladaptive reflexes^50–52^. Given its novelty, this short-latency effect should be regarded as the most provocative of our findings and treated as provisional until independently replicated.

This selective effect on corrective responses stands in apparent contrast to what taVNS did to unperturbed reaching, where its effects were global: endpoint variability was reduced and endpoints were biased toward the target center irrespective of target width, while movement velocity was unchanged. This global pattern cannot be read as a broad increase in neural gain, because the perturbation data show the opposite: corrective responses were not raised across targets but selectively suppressed. A more parsimonious account, invoking no second mechanism, links the global kinematic bias to the fixation requirement of our paradigm. Maintaining fixation on the central cross was task-relevant on every trial, independent of target width; if heightened arousal enhanced the processing of this uniformly relevant signal, it would reinforce the bias toward the central movement axis that sustained fixation already imposes, drawing endpoints toward the target center and reducing their spread for both targets. This interpretation is supported by our control experiment, in which the extended fixation period alone attenuated the functional distinction between narrow and wide targets (Fig. S.2). Crucially, the global kinematics and the selective reflex effects then reflect a single principle rather than two: arousal prioritizes processing according to task relevance, and whether this surfaces as a global or a target-specific change depends on the signal it acts on. Fixation was relevant on every trial regardless of target width, so enhancing it biased reaching globally, whereas a corrective response mattered only when a deviation threatened the goal, so suppressing it where it was dispensable separated wide from narrow.

An alternative account of the global kinematic effect appeals to the energetic cost of control rather than to fixation. A gain increase is metabolically costly and may therefore be deployed only where it is affordable: when no perturbation occurs, corrective demand is low and the increase can be expressed across both targets, producing the global kinematic effect; once a perturbation imposes a corrective demand, limited resources force prioritization, restricting the control to where accuracy is at stake and pruning the correction the wide target can tolerate. Resource-dependent prioritization of this kind fits proposed roles of noradrenaline in signaling and arbitrating the cost of action^20,53^. This account is also separable from the fixation one by experiment: removing the extended fixation requirement should leave the global kinematic effect intact under the energetic account but diminish it under the fixation account. Manipulating fixation directly would adjudicate between them, an issue for future study.

A pattern that cuts across both the kinematic and the reflex measures is that the effect of arousal depended on each participant’s initial behavior under sham. For endpoint eccentricity on wide target reaches, taVNS drew endpoints toward the center in participants who terminated far from it under sham and away from the center in those who already terminated near it, compressing the inter-individual distribution. The short-latency reflex on narrow-target reaches showed the same form of dependence: taVNS reduced responses in participants who showed large reflexes under sham and enhanced them in those who showed small ones. In both cases, arousal amplified what was weak and damped what was strong under sham. A cost-sensitive view of control offers a natural reading. Reaching to the center, like correcting a perturbation, is a more costly form of control than tolerating a deviation the goal can absorb, and participants differed in how much they exerted such control under sham. If arousal allocates control according to its cost, consistent with a proposed role of noradrenaline in arbitrating the cost of action^54^, it would raise it in those who exerted little and withhold it in those who exerted much, drawing each toward the level worth its cost rather than imposing a fixed, stereotyped change.

Several limitations temper our observations. The effects of taVNS were subtle: they did not alter the macroscopic kinematics of perturbed reaches, potentially because the amplified long-latency differentiation was carried by the difference index between targets rather than by either target response on its own. The short-latency suppression, though statistically clean and present in both agonist muscles, is a novel finding and warrants independent replication. Finally, targets were presented in separate blocks, so participants held sustained knowledge of the goal; whether the same goal-dependent modulation arises when targets are interleaved, and the goal must be resolved anew on each trial, remains to be established.

In conclusion, phasic boosts of arousal did not energize corrective control uniformly but reshaped it according to the goal of the movement, reducing corrective control for deviations the task could tolerate, as early as the short-latency reflex. These results bring the principle of selective gain, established in perception and memory, into the online control of movement, and recast arousal not as a global amplifier of the motor system but as a means of allocating corrective control to where it serves the task. More broadly, they place motor control within an emerging view of vagal and arousal signals as modulators that tune goal-directed behavior to its context rather than driving it indiscriminately^55^.

## MATERIALS AND METHODS

### 1. Participants, experimental setup and ethics statement

27 healthy participants (17 women, 24.0 ± 2.7 years old) were seated on an adjustable chair in front of a KINARM end-point robotic device (KINARM, Kingston, ON, Canada). Their head was tilted forward at approximately 45° from vertical with their forehead resting against a soft cushion attached to the frame of the workstation to ensure a good measure of the pupil size by the Eyelink 1000+ embedded in the KINARM device (see Fig. 1A). Participants were asked to keep their head as still as possible during any experimental block. The task required them to move a robotic handle with their right arm, keeping the elbow flexed at approximately 90° in the vertical plane between trials. The arm was hidden from view by visual blockers and a mirror which reflected a horizontal virtual reality display positioned above, showing the visual targets and representing the hand position as a white cursor (dot radius: 0.5 cm).

The KINARM device recorded the kinematics of the reaching movements. Surface electrodes (Bagnoli 119 EMG Sensor, Delsys Inc., Natick, MA, USA) measured the electromyography (EMG) activity of the Posterior Deltoid and Pectoralis Major, and pupillometry data were continuously recorded using the embedded Eyelink 1000+. All participants were right-handed, had normal or corrected to normal vision, no history of neurological or psychological disease and no physical injuries. They were all unaware of the purpose of the study. They signed an informed consent and received compensation for their participation. The study was approved by the Ethics Committee of the UCLouvain (reference: 2023/13JUL/322) and adhered to the Declaration of Helsinki.

### 2. Reaching task

Participants grabbed the handle of the robotic arm and performed reaching movements from a start target toward either a narrow (square) or a wide (rectangle) goal target. They were explicitly instructed that they could land anywhere within the goal target. Figure 1B illustrates the size and position of the start target along with narrow and wide goal targets superposed for visualization, although only one goal target was presented per trial. Figure 1C depicts the trial sequence, which always began with the start target at the bottom of the screen and the goal target at the top (again illustrated with both goal targets overlapping for schematic purposes). To initiate a trial, participants placed the cursor within the start target, which triggered the appearance of a cross at the center of this target. They were required to maintain gaze fixation on this cross during 9.5–10 s, after which the cross became a “+” giving them the movement go cue. Participants then had to reach toward the goal target within 0.3–1.2 s and stabilize the cursor inside it for 0.5 s to successfully complete the trial. The trials were concluded with performance feedback displayed for 2 s in the form of two thumbs: the left (black) thumb indicated whether the cursor had been stabilized in the goal target for the required 0.5 s, and the right (blue) thumb indicated whether the movement had been completed within the allotted time window (0.3–1.2 s). A “thumb up” signified success and a “thumb down” failure, with two thumbs up marking a fully successful trial. An intertrial interval of 3 s always followed before the next trial began. In all trials, a 4-s stimulation (taVNS or sham; see below) was delivered during the fixation phase. The stimulation was applied in the second part of the phase, starting 5.5 s after its onset and always ending before its offset. This ensured an additional 0–0.5 s of fixation after stimulation to avoid anticipation of the go cue. Movements initiated before the go cue led to immediate trial termination, and these trials, along with the subsequent one, were excluded during processing to ensure sufficient time elapsed between stimulations. In total, 20 trials out of 8424 trials were removed. Moreover, in two-third of the trials, a 9N step force (Fig. 1BC, dashed green line, noted as MP for “Mechanical Perturbation”) was applied randomly to the left or right once the hand crossed a horizontal positional threshold located 3 cm away from the start target. In these perturbation trials, participants were still required to reach the goal target and therefore needed to correct their movements.

### 3. Transcutaneous Vagus Nerve Stimulation

The setup and procedure for taVNS and sham stimulation followed the approach described in our previous work^28–30^ and are illustrated in Figure 1D. Stimulation pulses were produced with a Digitimer stimulator (Digitimer Ltd., model DS3), controlled by a Master-8 device (A.M.P.I., model Master-8). In the taVNS condition, electrodes were positioned on the left cymba conchae (anode) and tragus (cathode), areas densely innervated by the auricular branch of the vagus nerve^23,56,57^. This specific electrode montage has been shown to activate the nucleus tractus solitarius and, subsequently, the locus coeruleus^36^, a primary node of the ascending arousal system^14,17^. In the sham condition, electrodes were instead placed on the left earlobe, which is not expected to evoke brainstem or cortical activation^22,37,58^. Stimulation consisted of a 4s train stimulation of 200-µs pulses delivered at 25 Hz. To equate subjective sensation between taVNS and sham, intensity was individually calibrated for each participant and condition, following the procedure described by Su et al. (2025a; 2025b) and Denyer et al. (2026). Specifically, participants underwent a method-of-limits procedure to determine the taVNS intensity just below the pain threshold^36,59^. Current started at 0.1 mA and was increased by steps of 0.2 mA until participants rated the sensation as 9 on a 10-point scale (0 = no sensation, 10 = painful). This level was then confirmed in a descending series, in which intensity was decreased in 0.2 mA steps until ratings fell to 6 or lower. The final taVNS setting was defined as the average of intensities rated 8. For the sham condition, the same staircase was used, but participants judged the sensation relative to taVNS rather than giving numerical scores. The final sham intensity corresponded to the average level they perceived as matching the subjective intensity of taVNS at 8.

### 4. Experimental design

Participants attended a single laboratory session lasting approximately 3.5 hours, organized into three stages, as shown in Figure 1E. First, participants received training, which lasted until they felt comfortable with the reaching task and achieved an accuracy level of approximately 75–80%. Training blocks lasted about 5 min and were repeated 1 to 4 times depending on participants’ performance. Second, electrodes were placed on the left ear, and the stimulation calibration phase was conducted to determine the maximum comfortable intensity for both taVNS and sham conditions (see previous section). Finally, participants completed the main experiment, consisting of 8 blocks of 39 trials each (about 11 minutes per block). Within each block, 13 trials were unperturbed, 13 were perturbed leftward, and 13 rightward, in a random order. Target width (narrow vs. wide) alternated from block to block. Critically, the stimulation condition (taVNS vs. sham) alternated every other block: blocks were grouped into pairs that shared the same stimulation but differed in target width. Accordingly, the session comprised four block pairs with stimulation alternating across pairs, as for example: [taVNS: narrow, wide] > [sham: narrow, wide] > [taVNS: narrow, wide] > [sham: narrow, wide] (see Fig. 1E). The type of the first block (target width × stimulation) was counterbalanced across participants, yielding four possible block sequences.

In addition to these experimental blocks, EMG normalization blocks were interleaved every two blocks of trials, at the start of the session and at each change of stimulation condition, resulting in four normalization blocks in total (Fig. 1E). Each normalization block consisted of 16 trials during which only the start target was displayed. Participants were required to maintain the cursor within this area while countering a 9 N perturbation applied for 2.5 s, either to the left or right (randomized). Each block therefore contained 8 force applications in each direction. These blocks were used to measure the EMG activity elicited in the agonist muscle: the right posterior deltoid during leftward perturbations and the right pectoralis major during rightward perturbations. For each trial, mean EMG activity was computed in the 1.5–2.5 s window following mechanical perturbation onset. Values were then averaged across similar trials within a block, and finally across the four normalization blocks of the session. This yielded a baseline value used to normalize EMG data in the subsequent analyses (see below).

### 5. Data processing and End-point measures

#### 5.1. Electromyography data

The EMG data were recorded at a sampling rate of 1kHz and amplified with a factor 1k. The preprocessing was done with custom Matlab scripts (MATLAB 201 2022a, Mathworks Inc. Natrick Ma, USA), which filtered the data with a 6th order Butterworth band-pass filter with cutoff frequencies [20,250] Hz. Data were then exported to be analyzed using Python, as for the kinematic data. EMG signals were redressed and then normalized by dividing the activity of each muscle by its baseline value, computed during the EMG normalization blocks (see Experimental Design section). As for the kinematic data, normalized EMG signals were aligned to the moment the cursor crossed a threshold set to 3 cm beyond the start area, corresponding to perturbation onset. Finally, mean EMG activities during unperturbed trials were subtracted to the EMG activity during perturbed trials.

Analyses focused on reflex activity when the muscle acted as an agonist: leftward perturbations for the posterior deltoid, and rightward perturbations for the pectoralis major. Following established conventions^26,60^, reflex activity was divided into two epochs: the short latency response (SL; 20–50 ms) and the long latency response (LL, 50–100 ms) after perturbation onset. Normalized EMG in these two epochs served as the primary EMG outcome measures: SL-reflex and LL-reflex. Moreover, to assess target effects on the reflex activity, we computed a delta for each epoch and each participant, defined as the difference in activity between both goal targets (narrow – wide): ΔSL-reflex and ΔLL-reflex.

The kinematic data were recorded at a sampling rate of 1kHz using the KINARM software Dexterit-E (version 3.10). The preprocessing was done with custom Matlab scripts (MATLAB 201 2022a, Mathworks Inc. Natrick Ma, USA), which filtered the data with a dual pass 4th order low-pass Butterworth filter with a cutoff frequency set to 20Hz. Data were then exported to be analyzed using Python 3.11 via Anaconda Navigator (Version 2.5.0, Anaconda Inc.).

#### 5.2. Kinematics Data

Kinematic analyses focused exclusively on unperturbed trials, since the mechanical perturbation was primarily designed to elicit reflex responses, which were examined separately (see next section). Two main kinematic signals were extracted. Lateral eccentricity, shown in Figure 3B (left panel), was defined as the deviation of the hand position along the x-axis, computed as the distance between the hand x-coordinate and the straight line connecting the center of the start target to the center of the goal target (grey dotted line in Fig. 1B and in Fig. 3B, left panel). Forward velocity corresponded to the hand velocity along the y-axis and is displayed in Figure 3B (right panel). These two kinematic signals were aligned to the same time point as the hand trajectories and were analyzed within a window extending from 200 ms before to 700 ms after this event.

From these two signals, we extracted two measures of interest. First, we considered the absolute value of the lateral eccentricity at the end of the movement, i.e., at the moment when the forward velocity dropped below 3 cm/s^27^. This Eccentricity_Final_ therefore corresponded to the distance between the cursor and the goal target center at the end of the movement. This measure was computed only on trials with successful stabilization in the goal target (no position error). Second, we extracted the peak value of the forward velocity. This Velocity_Peak_ was computed only on trials without speed errors, i.e., when participants completed the movement within the required time window (0.3–1.2 s). For both endpoint measures, outliers were excluded through two successive iterations, retaining only values within the mean ± 2.5 standard deviations (SD). On average, this procedure yielded 24.3 valid trials per participant (out of 26) for the Eccentricity_Final_ analysis, with up to 15 trials excluded for a single participant. This corresponded to a total of 658 trials for the sham–wide target condition, 654 for the sham–narrow target condition, 656 for the taVNS–wide target condition, and 656 for the taVNS–narrow target condition, out of a possible 702 trials per condition. For the Velocity_Peak_ analysis, an average of 25.26 valid trials per participant (out of 26) were retained, with a maximum of 7 trials rejected for a single participant. It yields to 678 trials for the sham–wide target condition, 681 for the sham–narrow target condition, 682 for the taVNS–wide target condition, and 687 for the taVNS–narrow target condition, out of 702.

#### 5.3. Pupillometry data

The pupillometry (on the left eye) was recorded at a sampling rate of 500 Hz, during the whole trial. The pre-processing of these data was done using a custom Python script. First, blinks were detected and removed using a linear interpolation over a time window encompassing the blink and extending 100 ms before and after. Data were then low-pass filtered using a 3Hz fourth-order Butterworth filter, and trials with more than 50% of interpolated data during the fixation phase (see Reaching Task) were discarded. Pupil size data were converted in z-scores computed across all conditions for each participant and aligned to the sham or taVNS stimulation onset. These pupil z-scores were then normalized relative to the mean z-score obtained during the last second of the intertrial interval, when no target was displayed; this value was subtracted from z-scores obtained throughout the trial (see Fig. 4, lower panel). From these signals, we first computed the Pupil_Baseline_, defined as the mean pupil size during a 1-s window preceding the stimulation onset (see Fig. 1C and 4AA’). This measure served to evaluate baseline pupil size at trial onset (before stimulation) but when the start and goal targets were already displayed. Then, in order to allow for comparisons between conditions and participants, we subtracted this Pupil_Baseline_ from the remaining signal to obtain a relative measure of pupil dilation starting from stimulation onset (see Fig. 4, upper panel). From this signal, we selected a 2-s window, corresponding to the 2 last seconds of stimulation (see Fig. 1C and 4B). This Pupil_Stimulation_ measure reflects the pupil dilation elicited during taVNS or sham stimulation. The pupil analyses were restricted to unperturbed trials and to trials with leftward perturbations, corresponding respectively to those used for kinematic and deltoid reflex analyses.

### 6. Statistical analyses

Statistical analyses were conducted in Jamovi (version 2.5.3) and concerned kinematic (Eccentricity_Final_ and Velocity_Peak_), EMG (SL-reflex and LL-reflex as well as ΔSL-reflex and ΔLL-reflex) and pupillometry data (Pupil_Baseline_ and Pupil_Stimulation_). These data were analyzed at a single trial level using linear mixed-effects models with participants included as random intercepts, except for the variance of Eccentricity_Final_, ΔSL-reflex and ΔLL-reflex measures, which were analyzed at the participant level with ANOVAs, t-tests or with Wilcoxon signed-rank test when data were not normally distributed. Stimulation-Type (taVNS, sham) and Target-Width (wide, narrow) were the most common fixed factors. Pupil analyses also involved the factor Perturbation-Type (no, leftward). Post-hoc tests with Tukey corrections were applied to explore significant interactions.

For the mixed model analyses, estimated marginal means and standard errors are reported in the text and figures, whereas for t-tests or ANOVA analyses, observed means and standard errors are presented and shown as violin plots in figures.

