## Supplementary Materials for "Phasic arousal sharpens, rather than amplifies, goal-dependent feedback control"

### 1. *Statistical analysis*

taVNS appeared to standardize behavior, specifically for $Eccentricity_{Final}$ and SL-reflex, by increasing values that were low under sham and decreasing those that were high under sham. To quantify this effect, we used linear least-squares regressions assessing the relationship between taVNS-related changes (defined as the difference between taVNS and sham values) and the value under sham.

To ensure that the observed relationships were not driven by mathematical dependence between variables, we performed surrogate analyses. Specifically, we generated 5000 surrogate pairs of x and y datasets by sampling from normal distributions with the same mean and standard deviation as the empirical sham (x) and taVNS (y) data, respectively (as in Park et al., 2020; Kunz et al., 2024). For each surrogate pair, we computed the correlation between ($y - x$; mimicking the taVNS effect) and $x$, and extracted the slope of the corresponding linear regression. Empirical regression slopes falling outside the surrogate slope distribution indicate that the observed relationship is unlikely to arise from mathematical coupling and instead reflects a true behavioral dependency.

To further test whether taVNS-related changes were bidirectional (i.e., positive for some participants and negative for others depending on sham values), we performed a second surrogate analysis. We generated 5000 surrogate datasets by sampling from a normal distribution centered on 0 with a standard deviation matched to the residuals of the empirical linear regression. For each surrogate dataset, we estimated the intercept (offset). If the empirical intercept fell outside the surrogate intercept distribution, it was considered significantly different from zero, indicating that the taVNS effect was bidirectional rather than a single monotonic dependency on sham values. Finally, Levene's tests were performed to assess whether variance differed between conditions.

### 2. *Control experiment*

Eight healthy participants performed a control experiment (without sham or taVNS stimulation) designed to assess whether the extended fixation duration used in the main experiment influenced reflex responses. The task was identical to that of the main experiment: participants performed reaching movements from a start target toward either a narrow (square) or a wide (rectangular) goal target. They were explicitly instructed that they could land anywhere within the goal target, and the size and position of both the start and goal targets were the same as in the main experiment.

For half of the trials, the sequence of events was identical to that of the main experiment, with participants required to maintain gaze fixation on the cross on the start target (fixation trials). In the other half, the fixation period was omitted (no fixation trials). Once participants entered the start target, the go cue appeared after a random delay between 0 and 0.5 s, and no instructions regarding gaze were provided. In both trial types, a 9N step force was applied in two-thirds on the trials, randomly to the left or right once the hand crossed a horizontal positional threshold located 3 cm away from the start area, as in the main experiment.

As for the main experiment, participants completed 8 blocks of 39 trials. Trial type (fixation vs. no fixation) was held constant for the first four consecutive blocks and then switched for the final four blocks, whereas target width (narrow vs. wide) alternated from block to block. The starting block condition ([wide: no fixation], [narrow: no fixation], [wide: fixation] or [narrow: fixation]) was counterbalanced across participants.

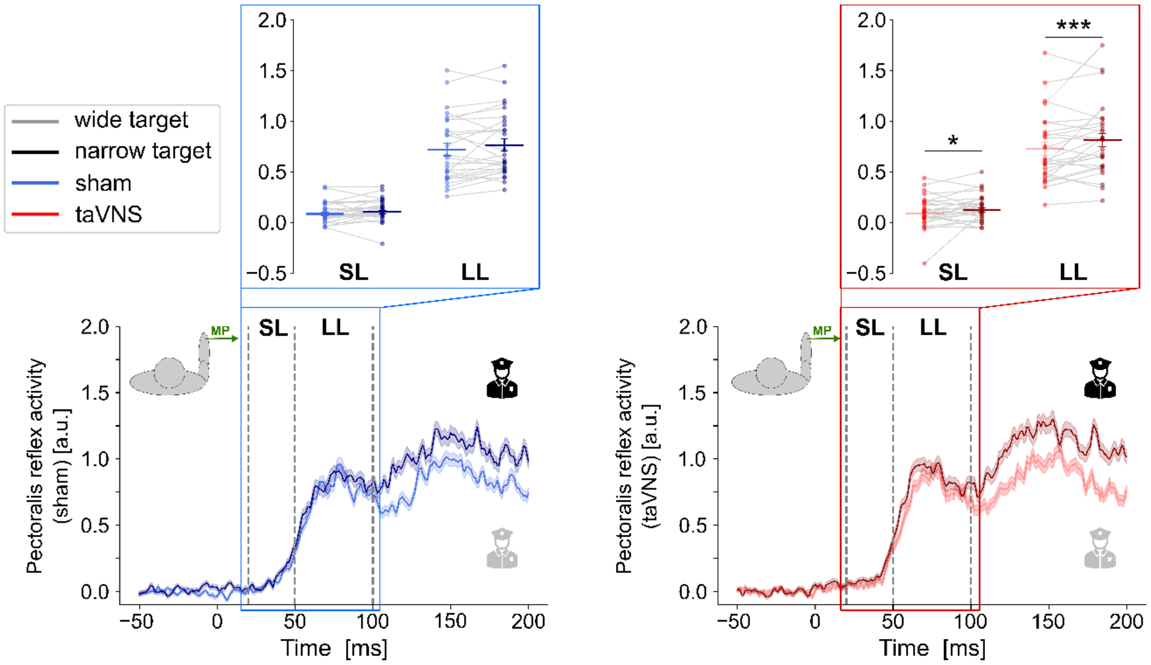


**Supplementary Figure S.1 – Pectoralis major activity with rightward mechanical perturbation (MP).** Data are shown for wide (light) and narrow (dark) targets under sham (blue) and taVNS (red) stimulation. **The lower panels show group-average EMG traces** (± SE) aligned to the mechanical perturbation onset and shown from −50 to 200 ms, under sham (left panel) and taVNS (right panel). Dashed vertical lines indicate the temporal boundaries of the short-latency (SL) and long-latency (LL) reflex epochs following the perturbation. **The upper panels show SL and LL reflex components.** EMM (horizontal lines) and EMSD (vertical error bars) are represented with individual participant data displayed as dots connected with thin grey lines, shown as a function of Target-Width, under sham (left panel) and taVNS (right panel). Note that, under sham, no statistical effect of Target-Width was observed on LL-reflex. *: $p < 0.5$. ***: $p < 0.001$. EMM = estimated marginal means, EMSD = estimated marginal standard deviations.

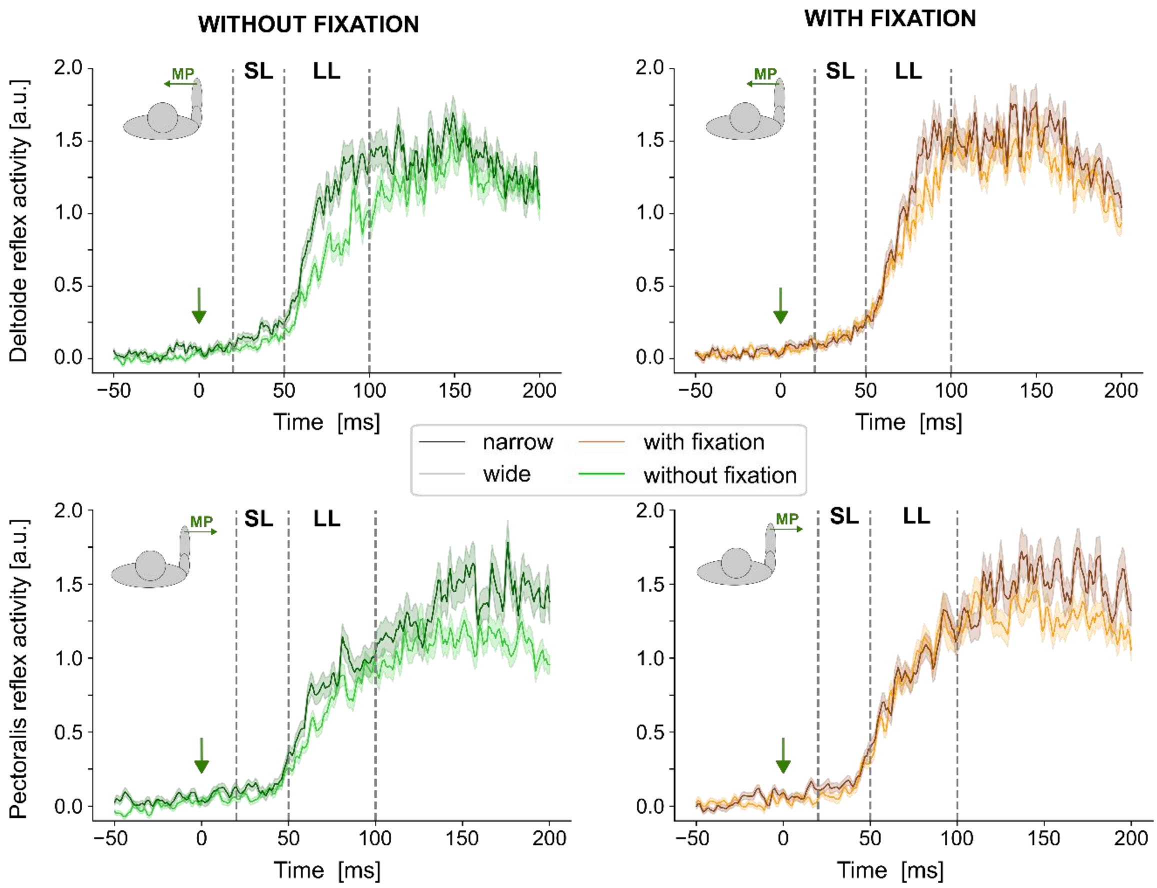


**Supplementary Figure S.2 – Results of the control experiment: group-average EMG traces for the posterior deltoid (upper panel) and the pectoralis major (lower panel).** Data are shown for wide (light) and narrow (dark) targets for trial with (orange) and without (green) the fixation period before the reaching movement. Mean (± SE) traces are aligned to the mechanical perturbation onset and shown from −50 to 200 ms, for trials without the fixation (left panel) and with the fixation (right panel). Dashed vertical lines indicate the temporal boundaries of the short-latency (SL) and long-latency (LL) reflex epochs following the perturbation.

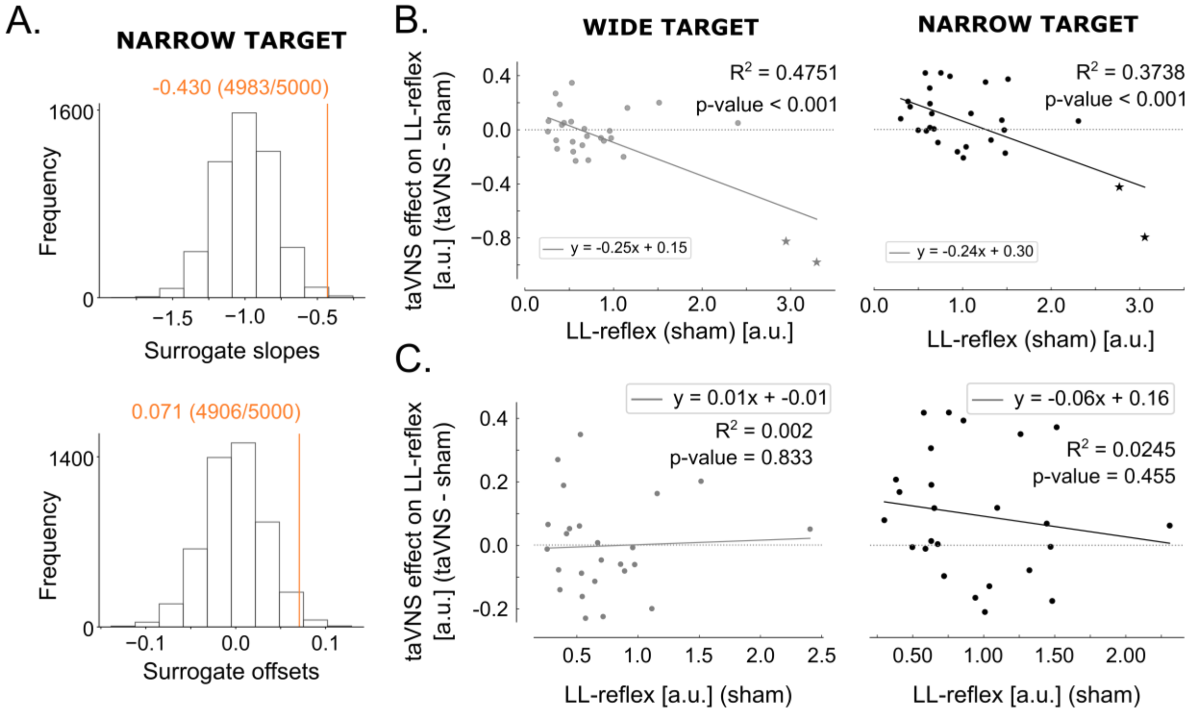


**Supplementary Figure S.3 – Additional analysis on posterior deltoid reflex activity. A. Surrogate analysis of SL-reflex.** Distribution of the surrogate slope (upped panel) and the surrogate offsets (lower panel) of SL-reflex for the narrow target. Empirical slopes and intercepts, obtained from the computed correlations, are indicated by orange vertical lines. Both the correlation slope (upper panel) and offset (lower panel) lay outside the surrogate distributions, indicating that the correlation on the right panel of Fig. 2C is not due to mathematical coupling and that the baseline-dependent taVNS effect is bidirectional. **B. Baseline-dependent taVNS modulation of LL-reflex.** Correlation between taVNS-related changes in LL-reflex (y-axis) and LL-reflex under sham (x-axis) for wide (left panel) and narrow (right panel) target. The two participants exhibiting exceptionally large sham LL-reflex values are represented as stars instead of dots. **C. Baseline-dependent taVNS modulation of LL-reflex without the two participants exhibiting large sham LL-reflex.** Correlation between taVNS-related changes in LL-reflex (y-axis) and LL-reflex under sham condition (x-axis) for the wide (left panel) and the narrow (right panel) target, without the two participants exhibiting exceptionally large LL-reflex amplitudes under sham conditions**.** Note that for both targets, the correlations were no longer statistically significant, indicating that the relationships observed in panel B were driven primarily by these two participants.

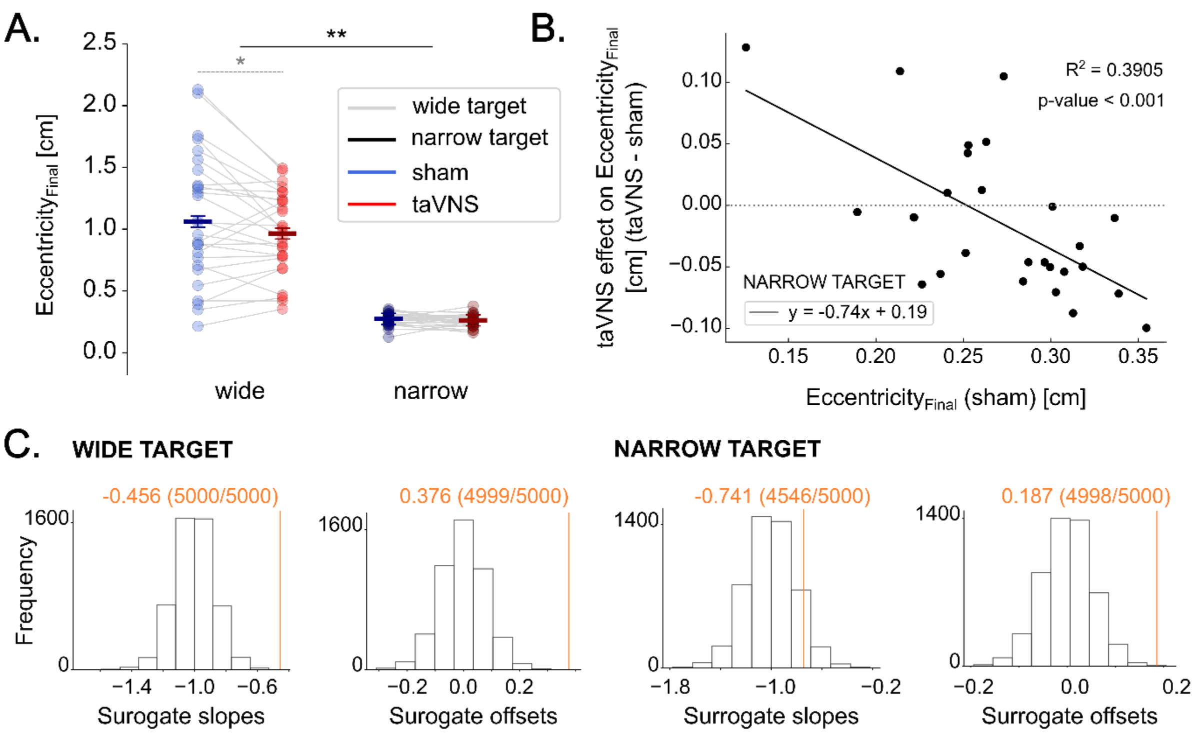


**Supplementary Figure S.4 – Additional analyses on Eccentricity$_{Final}$. A. Exploratory post-hoc analyses on Eccentricity$_{Final}$.** EMM (horizontal lines) and EMSD (vertical error bars) are represented with individual participant data displayed as dots connected by thin grey lines, shown as a function of Target-Width and Stimulation-Type. Note that this analysis suggests that Eccentricity$_{Final}$ was reduced under taVNS for the wide target, as shown by the dotted line. **B. Baseline-dependent taVNS modulation of Eccentricity$_{Final}$.** Negative correlation between taVNS-related changes in Eccentricity$_{Final}$ (y-axis) and Eccentricity$_{Final}$ under sham (x-axis) for the narrow target. **C. Surrogate analysis of Eccentricity$_{Final}$.** Distribution of the surrogate slope and the surrogate offsets of Eccentricity$_{Final}$ for the wide target (left panel) and the narrow target (right panel). Empirical slopes and intercepts, obtained from the computed correlations, are indicated by orange vertical lines. Note that for the wide target, both the correlation slope and offset lay outside the surrogate distributions, indicating that the correlation of Fig. 3D is not due to mathematical coupling and that the baseline-dependent taVNS effect is bidirectional. In contrast, for the narrow target, the slope fell within the surrogate distribution, indicating that the observed correlation can be attributed to mathematical coupling between the two variables. *: $p < 0.5$. **: $p < 0.01$. ***: $p < 0.001$. EMM = estimated marginal means, EMSD = estimated marginal standard deviations.
